# Environmental effects overtake selection to shape avian body size

**DOI:** 10.1101/2025.12.01.691188

**Authors:** Joshua Kenneth Robertson Tabh, Pierre Nyquist, Andreas Nord

## Abstract

Natural selection widely favours the largest within animal species. Why body sizes are declining in the world’s temperate birds is therefore mechanistically confusing and ecologically worrying. We argue that understanding the causes of these declines requires knowledge of how complete, body mass probability distributions – not just averages – are moving. Using data from 159 North American bird species – spanning 26 families, 53° of latitude, and 25 years – we show that body mass distributions are not only downgrading, but also narrowing, and losing their upper tails (the heaviest individuals), with the largest changes occurring at southern range limits. Combining fitness models with a novel, distributional Price equation revealed that these changes are increasingly driven by phenotypic plasticity, not selection or genetic drift, and may yet be reversible.

## Main text

> *“The Big Snake [Bashe, **巴蛇**] eats elephants and after three years it disgorges their bones. Gentlemen take a dose of this snake so that they will never have heart disease or illnesses of the belly.”*
>
> -Shanhai jing, chapter 10
>
> *“For by art is created that great Leviathan called a ‘commonwealth’ or ‘state’, which is just an artificial man—though bigger”*
>
> -Thomas Hobbes, Leviathan
>
> *“We’re not big, but we’re small.”*
>
> -Stuart MacLean, *in re* The Vinyl Cafe

Although a motley collection, this triple exergue reveals the almost salient connection between extreme size – be it large or small – and power and greatness in the human perception. While the reason for this connection is debatable, one might argue that its roots lie in evolutionary reality. Among animals, body size is almost always under positive selection, owing to its myriad competitive and reproductive advantages - so much so that its average within lineages tends to increase with evolutionary time [a phenomenon known as “Copes Rule”(1)]. By contrast, the sheer costs of a large size, including high resource demands and generally diminished annual reproduction rates (2), can lead small sizes to prevail in the game of fitness under certain conditions (3). But despite such evident links between size and fitness, our knowledge about how and why size varies at population-levels, and across space and near-time, is far from complete.

Since as early as the 20^th^ century, the average body size of many avian and mammalian species has diminished (4-6), with mean declines ranging between ∼1% per decade (5,7,8), and up to ∼12% per decade (9) in punctuated periods [albeit with considerable variability (10)]. That these declines coincide with increasing global temperatures has raised speculation about a climate-based cause [i.e. by increasing surface area-to-volume ratios, and thus, thermoregulatory costs (4,11)], and indeed, taxonomic “dwarfing” across earlier warming periods (e.g. the Palaeocene/Eocene Thermal Maximum, 55.8 mya) support this (12). We now know, however, that the causative picture is rich (13) and other drivers, including resource declines (14) – particularly for insectivorous species (15,16; but see 17) – and changes in species assemblage composition (18,19) are also at play. Yet even with this developing causative discourse, exactly how size declines have been mediated (i.e. via natural selection, phenotypic plasticity, or both) is hazy. Trends at northern latitudes imply relaxed selection against small body size as winters warm [the “warming winter” hypothesis (20-22); but see (23)], but direct changes in selection gradients, or genetic determinants of size, are scarce (24-26). Moreover, stunting effects of poor resource accessibility (9,27) and high temperatures during development (23,28) intimate phenotypic plasticity, and not selection, in driving size shifts. While both processes are likely operative, opinions on the relative contributions of each are divided.

We argue that a major barrier to understanding how size declines have occurred has been a failure to examine trends beyond central tendency. Longitudinal studies of mammalian and avian size have overwhelmingly focussed on changes to means (4-6), and less commonly, variance (29). Although informative, trends in central tendency and dispersion generally lack the diacrisis needed for critical hypothesis testing. For example, both relaxed selection against small body size and plastic reductions of realized body size equally result in declining means; so too should negative selection on large body size, whether from high thermoregulatory costs (11) or unattainably high energetic costs of maintaining such a size (Fig. 1). Distinctions in variance alone are also unclear. When viewed holistically, however, each of these mechanisms predict unique and testable differences on species-specific, body size probability distributions (Fig. 1). Should changes in whole body size distributions be clarified, we argue that the mechanisms driving size declines may therefore be inferred.

**Figure 1.**
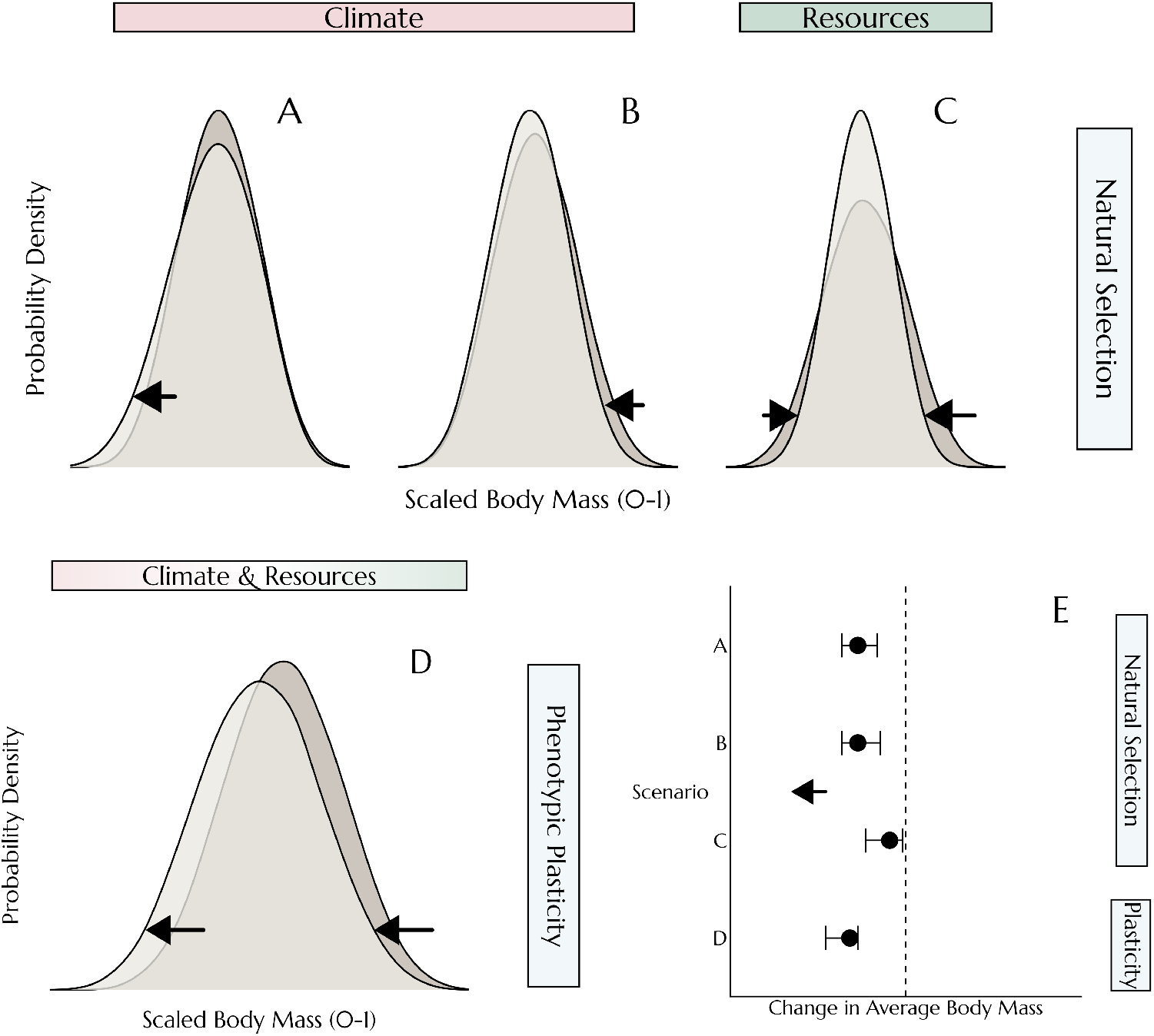
Outcomes of mechanistic hypotheses explaining changes in body size in response to climate warming (“Climate”) and resource limitation (“Resources”). Numerous hypotheses generate equivalent predictions about how body size averages should change. However, we show that predictions regarding changes in body size probability distributions vary. Panels **A** and **B** display predictions for two common hypotheses regarding how climate warming may diminish body size averages: (**A**) by relaxing mortality risk on small sizes^21^ or (**B**) by increasing selection against large body size, for example through increased thermoregulatory costs^11^. Panel **C** displays predicted outcomes of reduced resource availability on size distributions, specifically by imposing costs on both large sizes owing to high resource demands, and small size by limiting competitive ability, unequally but simultaneously. Last, panel **D** displays hypothetical outcomes for combined climate warming and resource limitation, acting through phenotypic plasticity. Here, climate warming truncates size at maturity through direct effects of higher developmental temperatures^47-48^, while reduced resources shift maximal attainable size downward. Differential outcomes of each hypothesis highlight the diacratic value of testing predictions at the distributional level. Panel **E** shows the similar change in averages from each hypothesis (or “scenario”) described in the previous panels.

Using a dataset documenting body mass of over 150 avian species across their North American ranges, we quantified how complete body mass distributions [here, representing body size distributions (30,31)] have changed in recent decades (Fig. 2A). Changes in distributions – particularly their midpoints, variance, skewness, and limits – are then used to infer the most likely mediator of widely-observed morphological changes. Our final dataset included 758,326 body mass measurements from 159 avian species (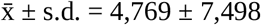 per species), 26 families (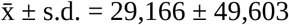 per family), and 26 years (1992-2018), collected during the reproductive period in summer. To more explicitly compare the contributions of natural selection and phenotype plasticity toward contemporary size shifts, we generated species-specific, body mass selection gradients (linking size with survivorship probability) from North American ringing records and temporal symmetry (Pradel) models, then partitioned the effects of these selection gradients on distributional changes using a novel adaptation of the classic Price equation (32). In its original form, the Price equation describes the degree to which a change in a species’ mean phenotype is explained by linear effects of selection and so-called “transmission” effects [which combine plasticity and drift (32)]. We adapted the Price equation into a ‘distributional’ analogue that partitions changes in a trait’s full probability distribution function between linear or non-linear selection gradients and complementary transmission effects. When paired with distributional comparisons, this adapted equation offers a more robust and flexible opportunity to pinpoint the precise determinants of phenotypic changes through time, such as avian size.

**Figure 2.**
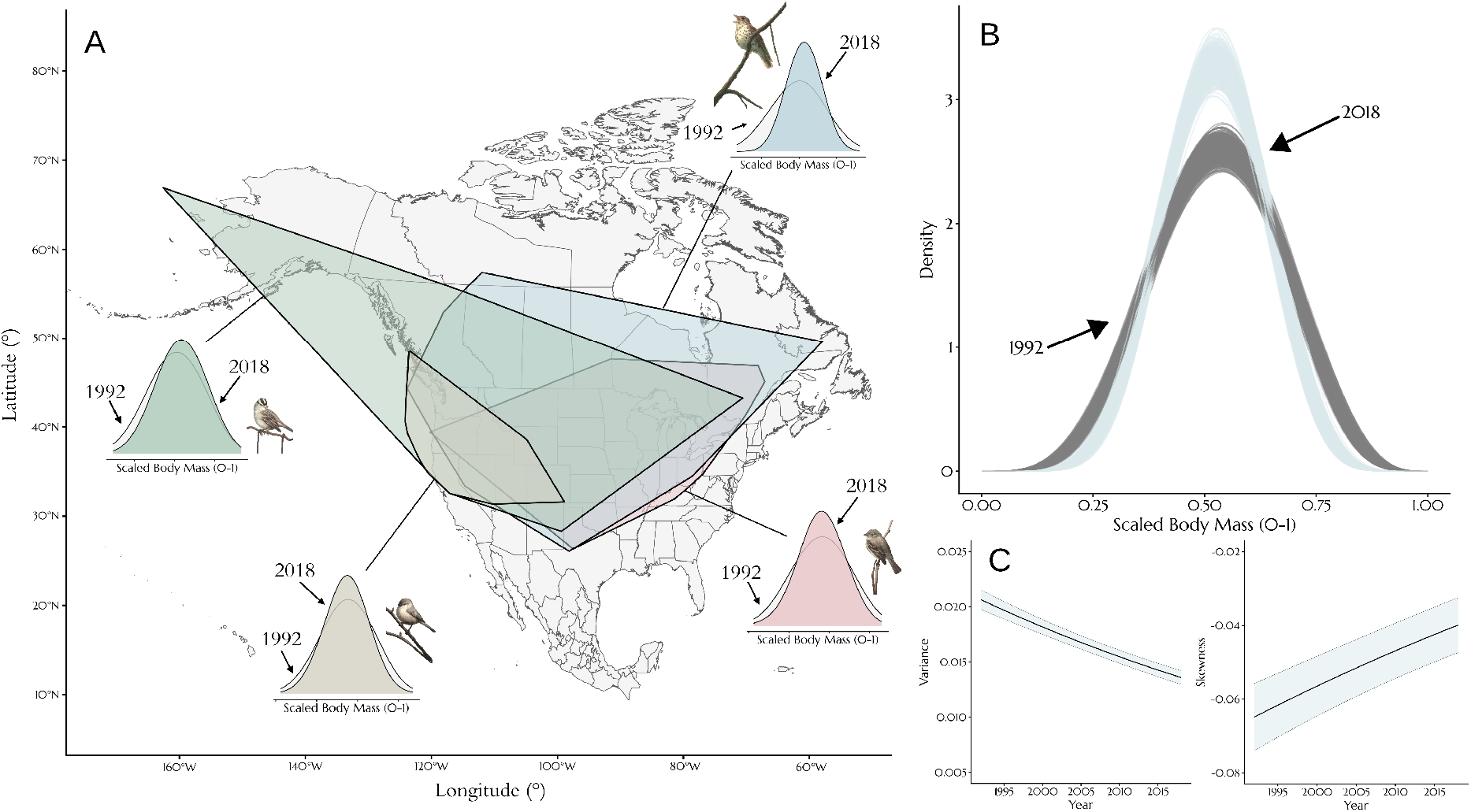
Declines in average size and size variance, and increases in size skewness, across 159 North American bird species. Body mass measurements were collected across observed species ranges and their distributions modelled across latitudes within and across ranges. A sample of species and distributional shifts at range medians is shown in panel **A** (complete set of distributional shifts in Supplementary Fig. 1). **B**. Estimated changes in average body size distributions between 1992 (grey) and 2018 (light blue) at median latitudes per species range, for all species in the data set. Lines indicate outcomes from distinct posterior samples drawn from a Bayesian mixed effects model. **C**. Changes in variance and skewness of body size distributions respectively, as estimates from outcomes of a Bayesian mixed effects model. Lines represent predicted trends, and bands (blue) represent standard deviations around predicted trends. Body size measurements are scaled between 0 and 1 per species and sex (where applicable).

### Body size distributions are contracting and shifting

Across all species and families assessed, body mass has predominantly declined (β = -0.012, or -1.2% per decade [posterior probability > 99.9%]), with average mass for 64.2% of species becoming smaller at their median latitudes. Percent changes in average mass ranged from -3.76% (the Black-throated Sparrow, *Amphispiza bilineata*) to +2.46% (the Carolina Wren, *Thryothorus ludovicianus*) per decade, aligning with previous estimates of size shifts in both North American and European songbirds (5,22). Coinciding with these declines, variation is body mass also decreased nearly ubiquitously (83.0% of species), averaging at a loss of 0.01 standard deviations (s.d. = 0.02) per decade (Figs. 2B-2C). This widespread and relatively rapid loss of mass variance in North American birds strongly contrasts the increasing trend in global, terrestrial mammals (29), and points to a disparity of mechanisms driving body size changes across animal groups. Whether this attrition of variance in birds is underpinned by a broader decline in genetic variation (33,34) is not clear, but holds substantial implications for long-term, species adaptability (35). Answering this question should therefore be an urgent priority.

Beyond shifts in central tenancy and dispersion, changes in the skewness of a phenotype can reveal crucial information about whether an adaptive optimum is moving, and how that movement is occurring (36-38). We found that body mass distributions have also become increasingly centralised, with historic left-skewness contracting in the majority of species (69.8%; Fig. 2C) owing to disproportionate losses above the mean (-0.17% of total probability density /decade). When viewed alongside changing means and variance, this illuminates a broad-scale downgrading – or left-shifting – of body mass distributions, not from an increasing presence of small body sizes, but from a decreasing presence of large body sizes. Supporting this, maximum observed body mass across species shrunk at a rate comparable to average body size (0.43%/decade; s.d. = 0.352; β = -0.011 [91.5%]; Fig. 3A), while minimum observed body mass remained unchanged across our 26 year observation period (β = 0.01 [23.1%]; Fig. 3B). These findings demonstrate that relaxed mortality risk on small size – stemming from warming winters [the warming winter hypothesis (20-22)] – is insufficient to explain the size declines observed across extant avian species. If distributional shifts were indeed driven by relaxed selection against small individuals, we would instead expect maximum body size to remain stable and minimum body size to diminish, the precise opposite of our observations.

**Figure 3.**
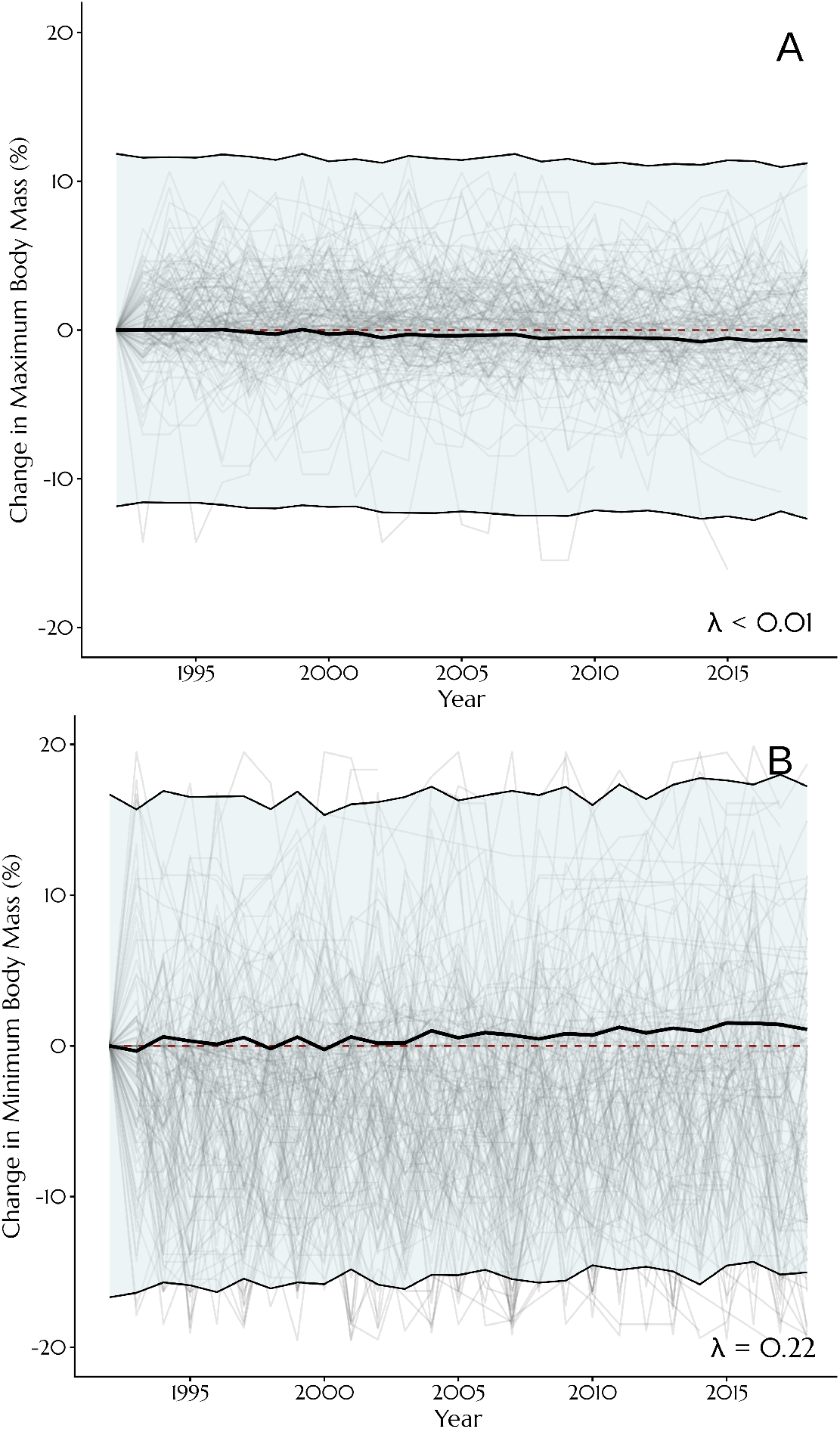
Declines in maximal body size, and stability of minimum body size across 159 North American bird species. Maximum (**A**) and minimum (**B**) body sizes represent maximum and minimum observed body mass, per species, and were natural-log transformed before use in Bayesian mixed effects model. Relative changes from baseline (1992) are displayed for adequate cross-species comparison. Grey lines represent single-species trends, while black lines indicate model fits. Bands (blue) display ± 1 standard deviation around trend lines. Red dashed lines display the 0-change line. In all panels, λ represents Pagel’s lambda.

To understand the generality of shifts in mass distributions, and better dissect their proximate mediators, we asked how they might have differed across the North American continent. We found that how body mass distributions have changed in recent years depended strongly on space and species-specific characteristics. Although shifts in average mass and mass variance did not differ across latitudes (Fig. 4; mean: β = -0.006 [46.4%]; variance: β ≈ 0 [6.2%]), shifts in variance did vary by relative latitude within species’ ranges (Fig. 4; mean: β = 0.008 [73.1%]; variance: β = 0.001 [82.5%]), indicating a possibly common, ultimate driver with intensity dependent on species’ characteristics (e.g. thermal tolerance or ecological niche) and not location itself. At southern-most observation points, variance declined at rates over-doubling those at northern-most observation points, reaching average loss rates as high as 13.3% per decade (relative to 6.1%) while changes in skewness remained unchanged (β ≈ 0 [5.6%]).

**Figure 4.**
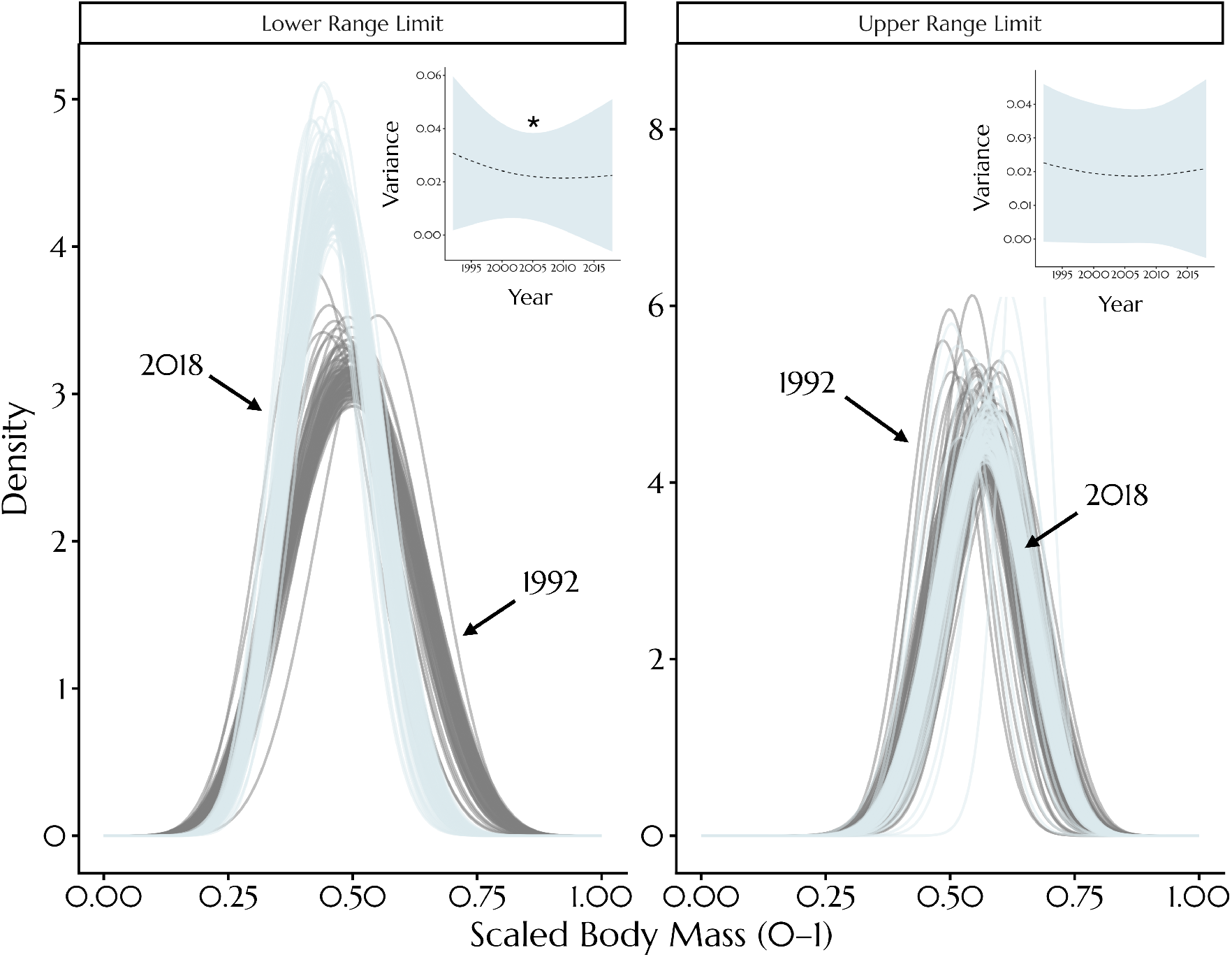
Declines in body size variance differ across species ranges. Temporal declines in body mass variance correlate with latitude within, but not across, species (β = 0.001 [82.5%]), with southern range limits experiencing the greatest losses of variance. Declines in average body mass are also greater at southern range limits, but with greater uncertainty (latitudinal effect: β = 0.008 [73.1%]). Lines in the large panels represent predicted body mass distributions across species, at southern-most and northern-most observation points (for each species individually). Blue lines indicate estimates for 2018 and grey lines indicate estimates for 1992; each individual line represents a posterior draw from a Bayesian beta-distributional model. Average changes in variance across time and species for each region (range limit) are shown in small insets. The asterisk indicates a trend with over 80% certainty.

While it is possible that northward movement from southern populations (39) has depleted some body mass variance at southern range limits, such movement should also yield concurrent increases in variance at northern range limits [owing from an influx of comparatively small individuals from southern populations (40)]. A lack of this paired signal indicates that poleward movement – the driver of taxonomic dwarfing in early, prehistoric warming periods (12) – is not sufficient to explain our observed shifts in size distributions through near-time. This is particularly significant, as it reveals that the mechanisms governing contemporary size changes in birds are not consistent with those governing size changes in historic eras. Instead, the comparatively faster declines in variance at southern range limits raise increased prevalence of extreme heat *per se* – highest at these locations – as a probable, common driver of distributional shifts, as already evidenced in some mammalian (41,42) and avian (23,28) populations. Still, given that changes in land-use tend to be most severe at lower latitudes [e.g. (43)], we cannot rule out the possibility of more rapid habitat and resource losses at southern range limits speeding local, distributional contractions. However, the similarity of distributional changes between low and high latitude species (indicated by the lack of a latitudinal effect across species; mean: β = -0.006 [46.4%]; variance: β ≈ 0 [6.2%]) indicates that this is likely insufficient to explain trends within species ranges.

### Environmental effects are increasingly shaping body size distributions

Effects of body mass on annual survivorship probability were predominantly quadratic across species and implicated stabilising selection as the dominant selective process shaping body mass distributions through time (mass^2^: β = -0.150 [95.3%]; mass: β = -0.009 [23.2%], mass^3^: β = -0.117 [13.7%]; controlling for variations in capture effort). Our analyses show that the intensity of this stabilising selection – and thus, selection against body mass extremes – has weakened (mass^2^ × decade: β = 0.218 [96.5%]), although considerable variability exists across species (54% of species with non-zero interaction terms at ≥80% posterior probability). If changes in selection gradients are mediating our changes in body mass distributions, this weakening selection against body mass extremes implies that mass variance should be increasing, not decreasing, as we observed (Figs. 2B-2C). This stark mismatch led us to evaluate the precise contributions of natural selection toward distributional shifts over time, using a novel Price equation (discussed above). Unlike the original Price equation, this revised equation offers two metrics of selective contribution toward a distributional change: (1) a “selective variance” (δ_2_S; ranging from 0-1), describing the ability of selection gradients to predict observed changes in probability distributions (similar to a Pearson R^2^), and (2) a “selective magnitude” (δ_1_S; also ranging from 0-1), describing the total magnitude of a distributional changes explained by selection gradients (representing a measure of strength or “penetrance”).

Our Price equation revealed that, since the beginning of our study period, selection gradients have become less able to predict annual changes in body mass distributions (selective variance: β = -0.55 [>99.9%]; Fig. 5A). Moreover, the strength of any detected selective effects on distributional changes has waned significantly (selective magnitude: β = -0.10 [>99.9%]; Fig. 5B). Phylogenetic effects of both changes were limited (Pagel’s λ = 0.04 [s.d. = 0.02] for each measure). Modelling the magnitude of selective effects linearly through time showed that since approximately 1990, non-selective effects – including phenotypic plasticity and genetic drift – had surpassed these selective effects as the largest determinant of body size distributions within species (Fig. 5B; averaged across species). Given many North American bird species have experienced large population declines in recent decades (44), we wondered if this rising prominence of non-selective effects could merely be explained by strengthening impacts of genetic drift in recent years [an expected outcome of decreasing effective population sizes (45)]. To test this, we compared changes in selective magnitude between species with increasing and decreasing population sizes across our last half-decade of observation (2010-2015), when the prominence of non-selective effects was clear (Figs. 5A-5B). Declines in selective magnitude were remarkably similar in both species groupings (Fig. 5C). Thus, rising non-selective effects are more likely explained by an increasing prominence of phenotypic plasticity, and not by increases in genetic drift.

**Figure 5.**
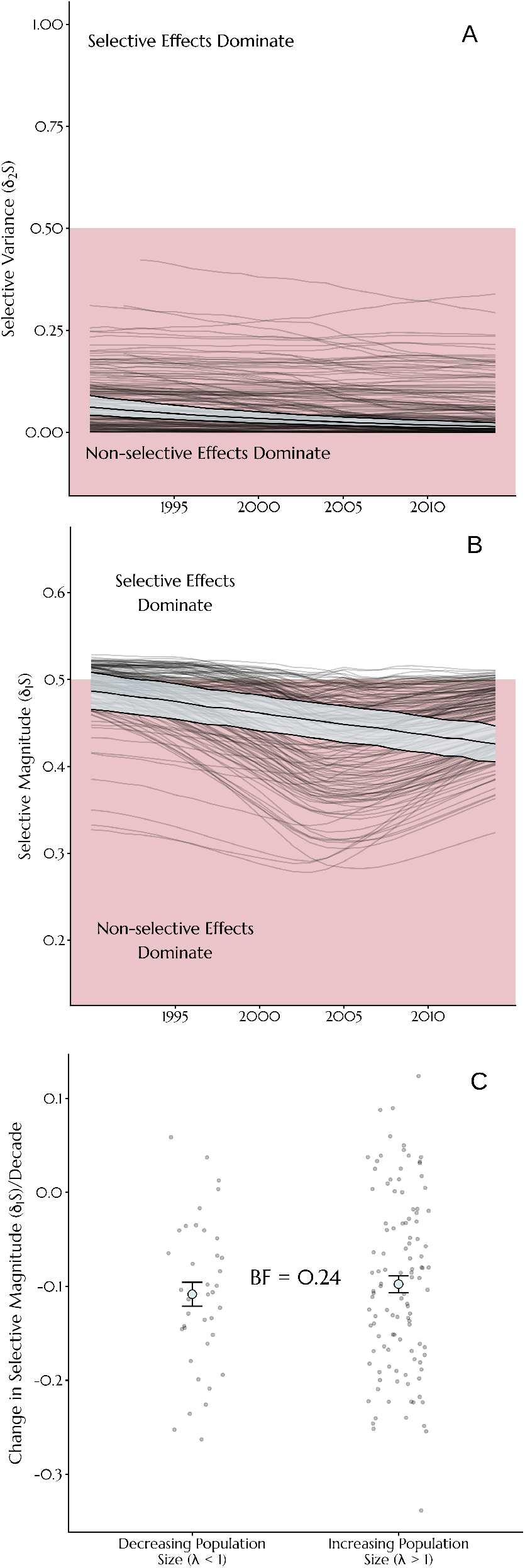
Phenotypic plasticity is increasingly defining body size distributions in North American birds (151 species). (**A**) Selective variance (δ_2_S) – derived from a distributional variant of the Price equation – describes the ability of a species’ body size selection gradient to explain changes in its body size distribution (similar to a Pearson R^2^). Values range between 0 and 1, with values above 0.5 indicating that natural selection is better able to explain distributional shifts than non-selective effects. (**B**) Selective magnitude (δ_1_S) describes the total extent to which estimated selective effects contribute to observed changes in body size distributions. A value above 0.5 indicate that selective effects explain more than half of the total distributional movement. Grey lines in **A** and **B** display trends for individual species. Black lines and blue bands display linear trend lines ± 1 standard deviation respectively, derived from Bayesian mixed effects models. **C**. Effect of population change (growth or decline) on linear changes in selective magnitude through time (2010-2015). Grey dots display values per species, blue dots display means, and errorbars display ± 1 standard error above and below means. Selection gradients and estimates of population change are derived from a temporal symmetry (“Pradel”) capture recapture model.

### Implications

Our results are consistent with temperature-mediated, phenotypic plasticity driving morphometric changes in North American birds. These findings confirm a generalised mechanistic basis across a suite of avian species (>150), with explicit support from an expanded and distributional Price equation. By lacking evidence for a selective basis (Fig. 5), we strongly question whether size shifts seen in these species may be considered outcomes of evolution [as also doubted on physiological grounds (13,31)].

Exactly how plasticity is operating to alter size distributions in North American birds is unclear. While increasing developmental temperatures – particularly at southern range limits [but see (46)] – may be directly truncating body sizes of young across their growth periods [reviewed in (47-48)], this mechanism is unable to explain the limited reduction of body size probability below medians (Fig. 2B). Alternatively, rising ambient temperatures may limit resources available for growth, either directly by changing the distribution and abundance of prey species (21), or indirectly by limiting offspring provisioning or viable foraging time (41,49). In this way, climate-mediated resource limitation may well explain our observed reduction in realised, maximal body sizes (Fig. 3A), as well as the slight loss of body size probability below medians if shortened foraging times motivate a need for increased, internal resource storage [i.e. as fat (50)]. Regardless, the reduction in body size and its variance reported here are likely to have meaningful consequences on environmental resilience and future persistence (1). Understanding whether plastic shifts in size distributions are offering hitherto unknown functional benefits will be a crucial next step toward predicting such consequences.

On a more fundamental level, our Price equation revealed a striking trend – that the relative contribution of natural selection toward avian body size has declined in recent decades. This trend raises a much deeper question about the role of environmental context (e.g. climate) in guiding the mechanistic determinants of animal phenotypes. Clarifying such a role is likely to influence our understanding of evolutionary histories, and is therefore deserving of further study.

## Acknowledgements

We thank Cecilia Lundberg, Jan-Åke Nilsson, Fredrik Andreasson, Elin Persson, Martin Stjerman and Erik Svensson for their useful insights during the construction of this manuscript.

## Funding

Wenner-Gren foundation grant UPD2021-0038 (JKRT, AN)

Sven och Lily Lawski foundation grant 20240523 (JKRT)

Stiftelsen Lunds Djurskyddsfond grant (JKRT)

Lars Hierta Minne scholarship 20221121 (JKRT). Funding for P.N. was provided by the

Swedish Research Council grant 2018-07050 (PN)

Swedish Research Council grant 2023-03484 (PN)

Swedish Research Council grant 2020-04686 (AN)

Swedish Research Council grant 2024-05362 (AN)

## Author contributions

Conceptualization: JKRT, AN

Methodology: JKRT, PN

Investigation: JKRT, PN, AN,

Visualization: JKRT

Funding acquisition: JKRT, PN, AN

Project administration: JKRT, PN, AN

Supervision: AN

Writing – original draft: JKRT

Writing – review & editing: JKRT, PN, AN

## Data, code, and materials availability

All data analyzed in this study are freely available through the MAPS program (https://www.birdpop.org/pages/maps.php) and the United States Geological Survey (USGS) Bird Banding Lab (https://www.usgs.gov/labs/bird-banding-laboratory/data). Collated versions are available in a private repository for review, available at the following links: https://figshare.com/s/3c5bbd0a89b4001ef87a, https://figshare.com/s/4d4f33022d54ba0709b2. All statistical code used in this study are also available on a private repository, accessible at the following links: https://figshare.com/s/4493c8f6d543adcea856, https://figshare.com/s/08baf3053e5d00f165be, https://figshare.com/s/961eb45b12d800827167.

## Notes

### Competing Interest Statement

The authors have declared no competing interest.

### Summary of Updates

(1) Manuscript structure. Results of the manuscript remain unchanged. Instead, the structure of the manuscript has changed to improve concision. (2) Figures. Figures in the manuscript have been updated for clarity.

