## Supplementary Information for "Environmental effects overtake selection to shape avian body size"

### Supplementary Materials

#### Materials and Methods

*Body mass data*

Body mass data for North American birds were obtained from the Monitoring Avian Productivity and Survivorship (MAPS) program (1), as organised by the Institute for Bird Populations (Petaluma, California). The MAPS program is a collaborative program that combines avian morphometric data from over 400 species captured across 1265 sampling locations, between the years of 1992 and 2018. Sampling locations in these years range from a minimum latitude of 26.065° to a maximum latitude of 69.382°, spread from the west to east of the North American continent [-165.669° to -57.606°; see (2)]. To reduce the influence of growth and migratory movement on body mass measurements – and thus the generality of our results – only MAPS data pertaining to adults captured between June and August inclusive were collected. Further, all body mass values exceeding species' medians ± 4 median absolute deviations were removed to reduce bias from species misidentification or measurement error.

Species inclusion was based solely on the sufficiency of measurements to estimate time-specific body mass distributions. As such, we excluded (1) all measurement years for a species where < 20 individuals were captured, and (2) all species where less than 4 measurement years remained after filtering by criteria 1. In total, our analysis included 758,326 body mass measurements from across 159 species ($\bar{x}$ ± s.d. = 4,769 ± 7,498 per species), 26 families ($\bar{x}$ ± s.d. = 29,166 ± 49,603 per family), and 26 years (1992-2018).

*Modelling shifts in species body mass across time*

Phenotype distributions in nature can vary in skewness, modality (i.e. unimodal vs. bimodal) and higher moments, as well as central tendency and dispersion (3-5). The beta probability density function ("pdf") is a flexible pdf able to capture variation in each for these metrics by adjusting the values of two shape parameters, α and β (or alternatively: μ, the distributional mean, and Φ, a scaling parameter). To model changes in body mass distributions, we therefore generalised these distributions per species using a beta pdf, with body mass first scaled between > 0 and < 1 (specifically, between $1.0\times10^{-5}$ – 1 - $1.0\times10^{-5}$) per species. To adjust for possible sexual size dimorphism within species, scaling was conducted within sex whenever sex was reliably determinable at capture (>75% of captures).

To estimate temporal changes in species' body mass distributions, we used a Bayesian mixed effects model with scaled body mass as the beta-distributed response variable, and with both μ (on the logit scale) and Φ (on the natural log scale) varying by model predictors. Sampling year was included as a continuous, population-level predictor (or "fixed effect"), and was median-centred and scaled by a factor of ten to ease model fitting. Given that body mass of many species varies by latitude (6), both a species' median latitude, and an individual's relative latitude for its species (in degrees latitude from the species' median) were included as additional, population-level predictors, with each parameters interacting with year to account for recent changes in species' ranges [e.g. (7)]. Last, to estimate intraspecific distributional changes, species identity was included as a group-level predictors of μ and Φ, with effects of year and relative latitude on μ and Φ also allowed to vary by species. The role of phylogenetic covariance on shaping distributional changes was evaluated in subsequent analyses (see below). Priors for our distributional model were informed by the findings of others (2,6,8-10) and by preliminary data visualisations. For population-level intercepts on μ and Φ, we used normally-distributed priors with means of 0.15 and 2.7 respectively, and standard deviations of 0.25, representing expected parameter estimates from cross-species histograms. For the effects of latitude and relative latitude within species, priors were also normal and assumed increases in distributional means with each variable (μ: latitude ~ $\mathcal{N}$[mean = 0.05, s.d. = 0.1], relative latitude within species ~ $\mathcal{N}$[mean = 0.05, s.d. = 0.1]), but no concurrent change in variance across variables (Φ: latitude across and within species ~ $\mathcal{N}$[mean = 0, s.d. = 0.05]). For our interactions between year and either latitude or relative latitude within species, all priors were conservative and assumed no effect on distributional means or variance (μ: $\mathcal{N}$[mean = 0, s.d. = 0.2] for both terms; Φ: $\mathcal{N}$[mean = 0, s.d. = 0.1] for both terms). At the species-level, priors were exponentially-distributed and as follows: μ and Φ: variation in intercepts ~ exponential(λ = 5), variation in slopes by year ~ exponential(λ = 10), and variation in slopes by relative latitude ~ exponential(λ = 10).

Posteriors were estimated using Hamiltonian Monte Carlo (HMC) sampling, with 20,000 sample iterations – thinned by a factor of 40 – drawn across 4 chains. The first 10,000 iterations from each chain were discarded as warm-up. To ease model initiation, a preliminary model run was conducted using meanfield automatic differentiation variational inference [ADVI (11)], and approximate posterior medians used to inform initial parameter values of our HMC chains. For this initial run, we used an η value (a step-size weighting variable) of 1.0, and 100,000 sample iterations. Chain mixing of our final model was evaluated by calculating Gelman-Rubin statistics [$\hat{R}$; (12)] contrasting them to 1.0, while autocorrelation within chains was evaluated by inspecting effective sample size to sample size ratios ($N_{eff}/N$) per parameter. $\hat{R}$ values fell between 1.00 – 1.02 and mean $N_{eff}/N$ values equaled 0.93 (s.d. = 0.12). Posterior predictive fits were diagnosed visually.

*Deriving variance and skewness from species' mass distributions*

For each species and observation year, we calculated the variance ($\sigma^2$) and skewness ($\gamma_1$) of body mass distributions, as derived from our distributional model (described above). For comparative purposes, each characteristic was also calculated at the minimum, median, and maximum observed latitude for each species. To do so, posterior predictions of μ and Φ for each species and context (n = 1000 each) were drawn from our distributional model, converted to standard beta-distribution shape parameters α and β ($\alpha$ = μ × Φ; $\beta$ = [1- μ] × Φ), and loaded into standard forms equations as follows:

$$\sigma^2 = \frac{\alpha\beta}{(\alpha+\beta)^2 \cdot (\alpha+\beta+1)}, \quad (1)$$

and

$$\gamma_1 = \frac{2 \cdot (\beta-\alpha)\sqrt{(\alpha+\beta+1)}}{(\alpha+\beta+2) \cdot \sqrt{(\alpha\beta)}}. \quad (2)$$

Resultant $\sigma^2$ and $\gamma_1$ values were then averaged per predictive scenario.

*Modelling changes in minimum and maximum body size through time*

To further our understanding of how body size distributions have, or have not, shifted through time, we tested whether species' observed minimum and maximum size measurements (here, natural log-transformed body mass) have changed across the same observation period (1992-2018). To do so, we constructed two Bayesian linear mixed effects models, one with each species' observed, annual minimum body size as the response variable, and the other with each species' observed, annual maximum body size as the response variable. In both models, error was treated as Gaussian, and we included year as the sole population-level predictor (median-centred and scaled by 10 as above), and both species and year-by-species as group-level predictors (intercept and slope respectively). To correct for phylogenetic non-independence between species, we allowed our model intercept and effect of year to vary by the phylogenetic covariance between species. Phylogenetic covariation was calculated from a consensus tree constructed using a recent phylogeny of birds (13) via Beast2's TreeAnnotator [(14) burn-in = 25% and node heights representing medians] as described elsewhere (15). Priors for both models described above were liberal, weakly informative. For our model predicting minimum observed size, priors for our population-level intercept and effect of year were normal, with means of 2.35 and 0 respectively, and standard deviations of 1 and 0.5 respectively. For our model predicting maximum observed size, priors for these parameters were largely identical, however, the mean of our intercept prior was set to 3. For both models, priors for all group-level effects were exponential with lambda values of 0.5. Sampling for each model was conducted across 4 HMC chains, totalling 50,000 iterations per chain, thinned by a factor of 20 and with the first 10,000 iterations discarded as warmup. $\hat{R}$ values averaged at 1 (s.d. = 0.001), while $N_{eff}/N$ values averaged 0.660 (s.d. = 0.305) across both models.

*Estimating annual survivorship at the species- and individual-level*

*Capture-mark-recapture model overview*

We next sought to estimate how body mass relates to fitness across species (i.e. via body mass selection gradients), and whether this relationship – if any – has changed through time. This estimation is a necessary step toward testing whether changes in body mass selection gradients alone are sufficient to explain recent shifts in body mass distributions (see below). Here, we used annual survivorship probability as the metric of fitness in our selection gradients, which is both readily estimated from capture-mark-recapture records and aligns with hypotheses seeking to explain morphometric shifts in birds and mammals [see (16)].

To estimate annual survivorship probability ($\phi$) at the species level, we used a hierarchical and Bayesian temporal symmetry model [or "Pradel model" (17)], adjusted to account for the presence of transients who are unlikely to be detected within a sampling area twice [adapted from (18)]. Briefly, this model uses both forward and reverse time to simultaneously estimate $\phi$, and an annual "seniority" probability ($\gamma$; i.e. that an individual captured at time $t$ was present in the capture area at time $t-1$) from capture-mark-recapture records, with each parameter being contingent on a probability of detection [$p$; reviewed in (19)], and a probability of residency within the sampling area ($r$; specifically, the probability that an individual captured for the first time is a resident in the area – equivalent to $r^F$ in (18). In this model, rates of population change ($\lambda$) directly represent the ratio of $\phi$ to $\gamma$ (17). This relationship is particularly useful for our study, because prior estimates of $\lambda$ are often available, and reparameterising the model to directly sample $\lambda$ instead of $\gamma$ improves estimational precision of $\phi$.

One known shortcoming of Pradel models is the assumption that all individuals have equal probability of capture. This implies that capture effort is constant across both space and time in a sampled population – an assumption that is difficult to satisfy when records are collated across independently organised observation sites such as in the MAPS data set. To overcome this limitation, we explicitly estimated mean annual capture effort ($\chi$) per species and year (described in the section below), then accounted for effects of that effort on $p$.

*Generalising the Pradel model for variations in capture effort*

To extend the utility of the Pradel model, we estimated mean annual capture effort ($\chi$) per species and year, for later integration in models (see below). Because effort required for detection may differ between naive and previously captured individuals (20), we estimated $\chi$ as latent determinant of two partial effort metrics: (1) the total number of observation days in a given year, per species, weighted by the probability of detection at each observation site ($\chi_C$; a metric of first capture effort), and (2) the average number of possible observation days before detection, per individual of a species, weighted by the probability of capture across active sampling sites per year ($\chi_R$; a metric of recapture effort). More specifically, we assumed:

$$\chi_{C_{sp,t}} = \sum_{cs=1}^{n_{cs}} N_{days_{cs,t}} \cdot \omega_{cs \vee sp}, \tag{1}$$

$$\chi_{R_{sp,t}} = \frac{\sum_{i=1}^{n_{sp,t}} \sum_{cs=1}^{n_{cs}} N_{days_{cs,t}} \cdot \omega_{cs \vee sp,i}}{n_{sp,t}}, \tag{2}$$

and

$$\begin{aligned} \ln(\chi_C + 1) &\sim N(\beta_C \chi, \sigma_C) \\ \ln(\chi_R + 1) &\sim N(\beta_R \chi, \sigma_R)' \end{aligned} \tag{3}$$

where *sp* represents species, *i* represents individual identity, *t* represents year, *cs* represents a capture site, $n_{sp,t}$ represents the number of individuals for a given species captured in year *t,* $n_{cs}$ represents the number of capture stations used in our study, $Ndays_{cs,t}$ represents the number of active days at a given capture site and year (defined as the number of days where at least one individual of any species was observed) and $\omega_{cs|sp,i}$ or $\omega_{cs|sp}$ indicates the probability of capture at capture site *cs* given a combination of species *sp* and individual *i.*

To first derive $\omega_{cs,sp}$, we divided the number of species observations at a capture site *cs* by the total number of observations across sites for species *sp.* To next derive $\omega_{cs|sp,i}$, we integrated Gaussian kernel densities drawn across all known capture-recapture distances (in km) per species, with lower limits of 0 and upper limits of the straight-line distances between an individual's initial capture site and possible recapture sites. Kernel bandwidths were set to 0.5 km, and as such, we assumed:

$$\omega_{cs_{i,sp}} = \int_0^{d_i} \left[ \frac{1}{n_{d,sp}} \sum_{j=1}^{n_{d,sp}} \frac{1}{0.5 \cdot \sqrt{2\pi}} e^{\frac{-1}{2}\left(\frac{d_{sp} - d_{j,sp}}{0.5}\right)^2} \right] dd_{sp} \tag{4}$$

where $d_i$ indicates the Haversine distance (in km) between the previous capture site of individual *i* and capture site *cs*, and $n_{d,sp}$ indicates the number of known capture-recapture distances for species *sp*. For the calculation of both $\chi_C$ and $\chi_R$, possible capture days represented only those during which a given species was expected to be present at a given site, in a given year. Relevant ranges were set by first modelling the very first known capture dates (within a year) per species as a function of Bayesian, Gaussian processes with latitude and longitude as predictors (n basis functions = 5; kernel = $\sigma^2 \cdot \exp\left(\frac{-(x - x')^2}{2l^2}\right)$, priors = flat; ε = Gaussian). Observation days across sites that preceded the expected first capture date from these models (the posterior medians), or preceded the observed first capture date in that year for a species (whichever was earliest) were then excluded from our estimates of $\chi_C$ and $\chi_R$. Last possible observation days used in $\chi_C$ and $\chi_R$ calculation were set to the final observation date for a given species and site, across years.

Once $\chi_C$ and $\chi_R$ were calculated, latent and "true" effort values ($\chi$) per species *sp* and year *t* were first approximated using a closed, single-iteration Newton-Raphson as follows:

$$\nabla_1\left(\chi_{sp,t}\right)=\frac{-\beta_C n_{sp,t}\cdot\left(\chi_{C_{sp,t}}-\beta_C\chi_{sp,t}\right)}{\widetilde{\sigma_C^2}}-\frac{\beta_R n_{sp,t}\cdot\left(\chi_{R_{sp,t}}-\beta_R\chi_{sp,t}\right)}{\widetilde{\sigma_R^2}}+\chi_{sp,t}$$

$$\frac{d}{d\chi_{sp,t}}\nabla_1\left(\chi_{sp,t}\right)=\frac{\beta_C^2 n_{sp,t}}{\widetilde{\sigma_C^2}}+\frac{\beta_R^2 n_{sp,t}}{\widetilde{\sigma_R^2}}+1 \quad (5)$$

$$\chi_{sp,t}^{(i+1)}=\chi_{sp,t}^{(i)}-\frac{\nabla_1\left(\chi_{sp,t}\right)\chi_{sp,t}^{(i)}}{\frac{d}{d\chi}\nabla_1\left(\chi_{sp,t}\right)},i=1,2$$

$$\chi_{sp,t}=\widehat{\chi_{sp,t}}=\chi_{sp,t}^{(2)}$$

where $\widetilde{\sigma_C}$ and $\widetilde{\sigma_R}$ represent the measured variance of $\chi_C$ and $\chi_R$ receptively when the alternate effort values (i.e. $\chi_R$ and $\chi_C$) are near their means (±0.05). $\beta_C$ and $\beta_R$ were estimated using a simultaneous and multivariate Bayesian, linear model following relationships described in eq. 3, with $\sigma_C$ and $\sigma_R$ being set as $\widetilde{\sigma_C}$ and $\widetilde{\sigma_R}$ respectively, scaled by the square-root of $n_{sp,t}$ (thus, assuming increased precision when the number of captures was high); where $n_{sp,t}$ for a given year $t$ was less than 1, scaling was ignored. For this model, priors for $\beta_C$ and $\beta_R$ were broad and normal, with means and variance of 1. Final estimates of $\chi$ across species and years represented posterior medians from this model (run across 4 HMC chains, each with 5,000 warm-up iterations, 5,000 sampling iterations, and a thinning factor of 10), and positive correlations between $\chi$ and both $\chi_R$ and $\chi_C$ were confirmed by visualisation.

*Pradel model specifications and capture-mark-recapture data augmentation*

For our Pradel model, we included capture-mark-recapture data from 1,620,420 individuals across 151 species, covering 26 families, 1786 capture sites, and spanning 25 years of observation (from 1990-2015). Primary data were obtained from the MAPS programme (1) and supplemented with additional capture data from the United States Geological Survey (USGS) Bird Banding Laboratory database.

All vital parameters in our Pradel model were estimated from independent functions. First, probability of detection ($p$) was modelled as a direct and linear function of effort ($\chi$), with both average $p$ and effects of $\chi$ on $p$ allowed to vary across species. Time was excluded as a determinant of $p$ since we assumed that most temporal effects on $p$ would be explained by temporal variations in $\chi$. Second, rates of population change ($\lambda$) were assumed to vary stochastically across sampling years, with variation in $\lambda$ across years differing by species. Similar to λ, the probability of residency ($r$) was also assumed to vary species-specifically across sampling years. However, unlike λ, $r$ is influenced by which observation sites were active in a given year – a variable with strong temporal inertia that is not truly stochastic. For this reason, $r$ values across years were modelled as first-order random walks per species. Last, annual probability of survivorship ($\phi$) was modelled as a linear function of time (i.e. by year, scaled as described for our distributional model), with average $\phi$, and the effects of time on ϕ, varying by species. A linear effect of time on $\phi$ was chosen to ease cross-species comparisons in future analyses. Together, formulae predicting $\phi$, $\gamma$, $p$ and $r$ were therefore as follows:

$$p \sim g^{-1}\left(\left[\beta_0+v_{0sp}\right]+\left[\beta_1+v_{1sp}\right]\cdot\chi\right)$$
$$\lambda \sim 2\cdot g^{-1}\left(\beta_0+v_{0sp,year}\right)$$
$$r \sim g^{-1}\left(\left[\beta_0+v_{0sp}\right]+\left[\sigma_{rw}\sum_{year=2}^{year} v_{1sp}\right]\right) \qquad (3)$$
$$\phi \sim g^{-1}\left(\left[\beta_0+v_{0sp}\right]+\left[\beta_1+v_{1sp}\right]\cdot t\right)$$

where *sp* indicates species, $g^{-1}$ indicates the inverse logit function (fixing select parameters on the probability scale), *year* indicates the sampling year, with 1990 equaling 1, *t* indicates the median-centred and scaled year, $\sigma_{rw}$ indicates a cross-species, random-walk scaling factor, and $v_{sp}$ indicates adjustments to population-level coefficients for species *sp*, or, in the case of *r*, the species-specific random-walk coefficient. Inverse-logit predictors of $\lambda$ are multiplied by 2 to allow λ to range between realistic extremes: 0 (a complete population crash) and 2 (a population doubling).

Priors for all cross-species predictors of model parameters were broad and moderately informed by estimates of others (21). For our cross-species intercept of *p*, we used a normal prior with a mean of 0 and standard deviation of 1.25, while for our effect of $\chi$ on *p*, we used a skew-normal prior with an $\xi$ of 0.5, $\omega$ of 1.5, and α of 5 (thus assuming a positive value). For both group-level effects of species on *p*, and the relationship between $\chi$ on *p*, we used half-normal priors with means of 0.1 and 0.05 respectively, and standard deviations of 0.15. Next, for our intercepts of $\lambda$ and *r*, we again used normally-distributed priors with means of 0 and -1 respectively (thus, assuming no population change and relatively low probability of residency), and standard deviations of 0.1 and 1 respectively. Our group-level effect of species-year on λ, and our random walk scalar were both modelled with exponential priors, with scale parameters (Λ) of 10 and 7.5 respectively. Last, for our cross-species intercept of $\phi$ and effect of scaled year on $\phi$, we used normally-distributed priors with means of 0 (setting reference $\phi$ at approximately 0.5 on the probability scale), and standard deviations of 0.75 and 0.1 respectively, while group-level effects of species on each were modelled with half-normal priors with means of 0.1 and 0.01 respectively, and standard deviations of 0.2 and 0.15 respectively.

Initial values for the HMC chains (n= 4) in our Pradel model were extracted from posteriors of an initial model executed using meanfield ADVI (as described above). In total, our final Pradel model was run across 4 HMC chains, and using 20,000 sampling iterations, thinned by a factor of 40, with the first 10,000 discarded as warm-up. All $\hat{R}$ values from this model fell between 1.00 – 1.03, indicating adequate chain mixing, and effective sample size to sample size ratios averaged 0.98 (± s.d. = 0.10), indicating minimal within-chain autocorrelation.

*Deriving individual-level survivorship*

Across individuals, the probability of recovering a captured individual (*R*) in a given year *j* may be described as:

$$R_{j,sp}=r_{j,sp}\cdot\phi_{j,sp}\cdot\left(\prod_{t=t_0+1}^{j-1}\phi_{t,sp}\cdot\left(1-p_{t,sp}\right)\right)\cdot p_{j,sp} \qquad (4)$$

where $t_0$ indicates an individual's year of last capture, and $p_j$, $\phi_j$, and $r_j$ represent averages for an individual's species (*sp*) in year *j* (see ref. 62). Using Bayes' Rule, the probability that an individual survives the year of its first capture (defined as $\phi_{j,t1}$) in the absence of subsequent recaptures can therefore be expressed as:

$$\left(\phi_{t1}\middle|R'\right)_{j,sp}=\frac{\left(1-p_{j,sp}\cdot r_{j,sp}\right)\cdot\phi_{j,sp}}{\left(1-p_{j,sp}\cdot r_{j,sp}\right)\cdot\phi_{j,sp}+\left(1-\phi_{j,sp}\right)}\ . \tag{5}$$

For subsequent years (where residency in the sample area is therefore assumed to be confirmed), $\varphi$ in the absence of recaptures may then be generalised as:

$$\left(\phi_{tx}\middle|R'\right)_{j,sp}=\left(\phi_{t1}\middle|R'\right)_{j,sp}\cdot\left(\prod_{T=1}^{x}\frac{\phi_{j+T,sp}\cdot\left(1-p_{j+T,sp}\right)}{\phi_{j+T,sp}\cdot\left(1-p_{j+T,sp}\right)+\left(1-\phi_{j+T,sp}\right)}\right)x\in\{2,3...L\} \tag{6}$$

where $L$ indicates the maximum number of years remaining in an individual of species *sp*'s expected lifespan. After $L$ has been reached, $\phi$ is then assumed to be 0.

To estimate $\phi$ across all known individuals and species, we used eq. 6 while also assuming that any future recapture events for individual $i$ implies that $\phi = 1$ in year $j$. In this context, $p_{j,sp}$, $r_{j,sp}$ and $\phi_{j,sp}$ were predicted from our Pradel model (posterior medians), with capture effort – required to predict $p_{j,sp}$ – representing individual estimates in place of species-level averages, and $r_{j,sp}$ equaling 1 if an individual was previously captured at least once (confirming their residency status). To determine $L$, we calculated each species' apparent longevity [i.e. their maximum lifespan (22)] from capture-mark-recapture records as the maximum time between capture and recapture events plus the age at first capture (assumed to be 1 if unknown) within that species. $L$ for a given individual was then set to its species' apparent longevity less the number of years between an individual's estimated age from capture records (again, 1 if unknown).

*Constructing selection gradients*

Only observations where both $\phi$ and body mass were available in a given year were used to construct the selection gradient model. Final data used in this analysis thus included 85,880 observations collected between 1992 and 2014, and spanning across 150 species and 26 families.

To determine: (1) how body mass relates to fitness across species, and (2) how this relationship may have changed through time, we combined annual survivorship probabilities with scaled body mass measures to construct time-varying selection gradients, following standard approaches (23). Specifically, we constructed a Bayesian linear model with individual-level survivorship estimates ($\phi$; derived in the section above), divided by their species-specific means (23) as our response variable, and relative body mass (scaled between 0 and 1 within species and sexes, then centred at 0.5), year (scaled by a factor of 10 and median centred, as described above), and the interaction between relative body mass and year as population-level predictors. Hypotheses seeking to explain shifting morphology in birds often assume non-linear patterns of selection (Fig. 1). Thus, to allow for both linear and non-linear effects of body mass on $\phi$, we also included squared and cubic forms of relative body mass as predictors, with year interacting with each level. To account for possible spatial variations in slopes of $\phi$ by year, we also included latitude and longitude of an observation as a Gaussian process predictor (n = 3 basis functions; kernel = $\sigma^2\cdot\exp\left(\frac{-\left(x-x'\right)^2}{2l^2}\right)$. Finally, to consider possible variation in body mass, temporal, and spatial effects on $\phi$ by species, we included group-level intercepts of species on $\phi$, as well as group-level slopes of mass, time, the interaction between each, and space (our Gaussian process) per species (where Gaussian processes were constructed group-wise), assuming no correlation between each. We assumed a Student's *t*- distributed error for this model, owing to high excess kurtosis apparent in data visualisations. Priors for our model describing annual survivorship probability ($\phi$) were weak and conservative, assuming no effect of body mass and year on survivorship (intercept ~ $\mathcal{N}$[mean = 1, s.d. = 1], body mass ~ $\mathcal{N}$[mean = 0, s.d. = 4], body mass$^2$ ~ $\mathcal{N}$[mean = 0, s.d. = 16], body mass$^3$ ~ $\mathcal{N}$[mean = 0, s.d. = 16], year ~ $\mathcal{N}$[mean = 0, s.d. = 0.5], body mass × year $\mathcal{N}$[mean = 0, s.d. = 2.5], body mass$^2$ × year $\mathcal{N}$[mean = 0, s.d. = 2.5], body mass$^3$ × year $\mathcal{N}$[mean = 0, s.d. = 2.5], all group-level effects of species ~ exponential[Λ = 2.5], ℓ[Gaussian process length-scale parameter] ~ exponential[Λ = 0.5], standard deviation of Gaussian process ~ Gamma[α = 2, β = 0.75]). Error in this model was assumed to be near-Gaussian (ν ~ Gamma[α = 2, β = 1.25]). This model was run across 4 HMC chains, again using 14,000 sampling iterations thinned by a factor of 10, with the first 8,000 discarded as warm-up. $\hat{R}$ and Neff/N values for all coefficients fell between 1.00 – 1.01, and 0.55 - 1.08 respectively.

*Partitioning effects of selection and plasticity on changes in body mass distributions*

To partition the contributions of natural selection and phenotypic plasticity toward changes in species' body mass distributions, we constructed a novel, "distributional" Price equation that accepts phenotypic change functions on the left side of the equation, instead of changes in mean phenotypes. Briefly, the Price equation is an exact calculation that describes the change in the average value of a trait within a population or species. In its original form, this change is defined as the sum of the scaled covariance between that trait and relative fitness (a "selection term"), and the combined effects of plasticity, drift, and mutational effects [often defined as the "transmission", or "non-selective" term (24)]. This equation does not allow for partitioning of selective and non-selective effects toward more complex changes in phenotypes within a population, for example, in skewness and higher moments.  Moreover, by defining the selection term as covariance between two continuous variables, Price's equation does not allow for integration of, or distinguishment between, forms of "variance selection" (e.g. stabilising selection, disruptive selection), or otherwise non-linear selection. Our distributional equation accommodates both and offers novel opportunities to apply Price's identity to answer questions about the role of selection in shaping traits at a holistic level.

Our distributional Price equation is derived as follows: setting $z$ as a trait value, and $p_1$ and $p_2$ as probability density functions (pdfs) across the set of $z$ values ($Z$) at time-points 1 and 2 respectively. The change in a trait's pdf [$\Delta p(z)$] between two adjacent time-points is given as:

$$\Delta p(z) = p_2(z) - p_1(z) \tag{7}$$

or equivalently:

$$\Delta p(z) = \left(p'_1(z) - p_1(z)\right) + \left(p_2(z) - p'_1(z)\right) \tag{8}$$

where $p'_1(z)$ represents an intermediate pdf, and the first and second summands represent changes in $\Delta p(z)$ driven by selective (henceforth $\nabla_{NS}$) and non-selective effects ($\nabla_{EC}$) respectively [*sensu* (25)]. Setting $f_1$ as a selection gradient (i.e. a function relating a given trait value to its relative fitness value), and $E_{p_1}[f_1]$ as the expectation of $f_1$ under $p_1$, $p'_1(z)$ may be explicitly defined as:

$$p'_1(z) = \frac{f_1(z)\,p_1(z)}{E_{p_1}[f_1]} \tag{9}$$

and $\nabla_{NS}$ therefore as:

$$\begin{aligned}\nabla_{NS} &= \frac{f_1(z)\,p_1(z)}{\mathbb{E}_{p_1}[f_1]} - p_1(z) \\ &= \left(\frac{f_1(z)}{\mathbb{E}_{p_1}[f_1]} - 1\right) p_1(z)\end{aligned} \quad . \tag{10}$$

Next, using a displacement kernel ($K_D$) to describe the probability of a given trait value ($z$) being obtained from another trait value ($y$) within the same set $Z$ (i.e. via plasticity or imperfect inheritance), $\nabla_{EC}$ is defined as:

$$\nabla_{EC} = \int_{y \in Z} K_D(z|y)\, p'_1(y) - p'_1(z)\, dy\,, \tag{11}$$

or:

$$\nabla_{EC} = \int_{y \in Z} \left[K_D(z|y) - \delta(z-y)\right] p'_1(y)\, dy \tag{12}$$

where δ represents the Dirac delta (setting $p'_1(y) = \int p'_1(z)\,\delta(z-y)\,dy$). Combining eq. 10 and 12, and expanding $p'_1(y)$, we obtain the near-final form:

$$\Delta p(z) = \left(\frac{f_1(z)}{E_{p_1}[f_1]} - 1\right) p_1(z) + \frac{\int_{y \in Z} p_1(y) f_1(y)\left[K_D(z|y) - \delta(z-y)\right] dy}{E_{p_1}[f_1]}; \tag{13}$$

Given $K_D$ is often unknown, the non-selective term ($\nabla_{EC}$) is not always calculable and may only be solved given a known change in a probability distribution [$\Delta p(x)$] and a known selection gradient $f_1$. In these cases, setting the non-selective term to $\varepsilon_1$, eq. 13 is simplified to the more useful form:

$$\Delta p(z) = \left(\frac{f_1(z)}{E_{p_1}[f_1]} - 1\right) p_1(z) + \epsilon_1(z). \tag{14}$$

To evaluate and compare the relative contributions of both the selection and transmission term toward $\Delta p(x_2)$, we offer two approaches: one in $L_1$ form (analogous to regression slopes) and one in $L_2$ form (analogous to Pearson $R^2$ values). For the $L_1$ solution, contributions of the selective and non-selective terms (set as $\partial_1 S$ and $\partial_1 E$ respectively) are defined as:

$$\partial_1 S = 1 - \frac{\left(\int \left|\Delta p(z) - \left(\frac{f_1(z)}{E_{p_1}[f_1]} - 1\right) p_1(z)\right| dz\right)}{\int |\Delta p(z)|\, dz}; \partial_1 E = 1 - \partial_1 S\,, \tag{15}$$

or more stably as:

$$\partial_1 S = \frac{\int |\nabla_{NS}| dz}{\int |\nabla_{NS}| dz + \int |\nabla_{EC}| dz} \partial_1 E = 1 - \partial_1 S, \tag{16}$$

where $\nabla_{EC}$ is obtained from $\Delta p(z)$ - $\nabla_{NC}$. We define $\partial_1$S descriptively as "selective magnitude".

For the $L_2$ solution, these contributions (set as $\partial_2$S and $\partial_2$E respectively) are thus:

$$\partial_2 S = \frac{\left( \int \Delta p(z) \cdot \left( \left( \frac{f_1(z)}{E_{p_1}[f_1]} - 1 \right) p_1(z) \right) p_1(z) dz \right)^2}{\left( \int (\Delta p(z))^2 p_1(z) dz \right) \cdot \left( \int \left[ \left( \frac{f_1(z)}{E_{p_1}[f_1]} - 1 \right) p_1(z) \right]^2 p_1(z) dz \right)} ; \partial_2 E = 1 - \partial_2 S, \tag{17}$$

with $\partial_2$S being defined, descriptively, as "selective variance". Validation of our derivations is described in the following section.

*Validating Price derivations*

To validate our functional Price equation, we generated counter-factual body mass pdfs per species and year, where changes in these pdfs are explained purely by estimated selection gradients operative at that time. If our estimated contribution of selection toward a change in a mass pdf is high), the difference between this counter-factual mass pdf and an observed, "true" mass pdf for the same year and species should be small. To formally test this prediction, we first constructed counter-factual mass pdfs as follows:

$$h(x)_{sp,t}{}' = \frac{f(x-0.5)_{sp,t-1} \cdot Beta(x|\alpha_{sp,t-1}, \beta_{sp,t-1})}{\int_0^1 \left( f(x-0.5)_{sp,t-1} \cdot Beta(x|\alpha_{sp,t-1}, \beta_{sp,t-1}) \right) dx} \tag{6}$$

where *x* indicates a scaled body mass, *f(x)* again represents a selection gradient for a given species *sp* and year *t* (where *x* was centred at 0), *α* and *β* are posterior medians predicted per species and year at a species' median latitude (from our distributional model), and *h(x)'* represents the counter-factual mass pdf. To approximate the precise form of *h(x)'* – a requirement for KL divergence calculation – we used linear interpolation occurring across 250 evenly spaces points. KL divergences between *h(x)'* and "true" body mass pdfs (at equivalent time points and species) were then calculated using standard methods:

$$KL_{sp,t} = \int_0^1 h(x)'_{sp,t} \cdot \ln\left( \frac{h(x)_{sp,t}{}'}{Beta(x|\alpha_{sp,t}, \beta_{sp,t})} \right) dx \tag{7}$$

where *KL* represents the KL divergence for a given species *sp* and time *t*. KL divergence values were then plotted against $\partial_1$S from our functional Price equation at equivalent time-points per species to confirm a negative relationship. We then confirmed this negative relationship by modelling natural-log transformed KL divergence values as a linear function of $\partial_1 S$ using simple, Bayesian linear regression, with weak priors on the effect of either selective contribution term ($\mathcal{N}$[mean = 0, s.d. = 1]) and on model intercepts ($\mathcal{N}$[mean = -5, s.d. = 5]). This model was run using 4 HMC chains, sampled with 20,000 iterations, thinned by a factor of 20, and with the first 10,000 iterations removed as warm-up. $\hat{R}$ and $N_{eff}/N$ values for all parameters ranged between 0.99 and 1.01, and 0.93 and 1.01 and respectively.

Our selective magnitude term ($\partial_1 S$) negatively correlated with the KL divergence between true body mass distributions and expected (or "counterfactual") distributions, derived from selective effects alone ($\beta$ = -0.30 [82.8%]). More specifically, expected body mass distributions – solely defined by selective effects – were more similar to true body mass distributions when $\partial_1 S$ was high. This finding confirms that the selective contribution derived from our Price equation ($\nabla_{NS}$) indeed positively reflects that magnitude of estimated selective effects.

*Evaluating changes in the contribution of natural selection toward body mass distributions*

We calculated selective magnitude ($\partial_1 S$) and selective variance ($\partial_2 S$) for all species with defined body mass selection gradients (as described above; setting latitude and longitude as the geographical centre of each species' range, and extracting posterior medians) and time-specific body mass distributions (calculated previously, and assuming median latitudes per species) between the years of 1990 and 2015. Given that both $\partial_1 S$ and $\partial_2 S$ are direct solutions, functions of each across time have a precise polynomial form. However, we were interested in understanding the general, linear trajectories of these polynomial forms through time. As such, once calculated, we constructed two Bayesian linear models with logit-transformed $\partial_1 S$ and $\partial_2 S$ as Gaussian-distributed response variables, and with year (median-centred and scaled by decade) as the sole population-level predictor in both models. To understand the role of phylogeny in shaping temporal changes in $\partial_1 S$ and $\partial_2 S$, we allowed both intercepts and temporal slopes to vary by species and phylogenetic covariance between species, as group-level effects. Calculation of phylogenetic covariation is described earler. Priors for model intercepts we normal with means of 0 and standard deviations of 2 in natural log space. For effects of time on $\partial_1 S$ and $\partial_2 S$, we also assumed normally-distributed priors with means of 0 and standard deviations of 1. Last, for effects of species identity and phylogenetic covariance on both intercepts and time-dependent slopes, we used exponential priors with lambda values ($\Lambda$) liberally set to 0.5.

Models describing $\partial_1 S$ and $\partial_2 S$ across species and time were run using 20,000 sampling iterations across 4 HMC chains, each thinned by a factor of 20, and with the first 10,000 iterations in each chain being removed as warm-up. Similar to previous models, $N_{eff}/N$ values average > 0.8 across all parameters ($\partial_1 S$ model: s.d. = 0.05; $\partial_2 S$ model: s.d. = 0.09) and $\hat{R}$ values fell consistently between 0.99 and 1.01.

*Statistical reporting and overview*

All statistical models were run in R (version 4.4.1; R Core Team, 2025) using Stan (version 2.32.2), as operated through the R packages "cmdstanR" (26) and "brms" (27). Model coefficients represent posterior modes, unless otherwise stated. Posterior probabilities represent the smallest, symmetrical highest posterior density intervals around posterior modes that include 0.

### Figures

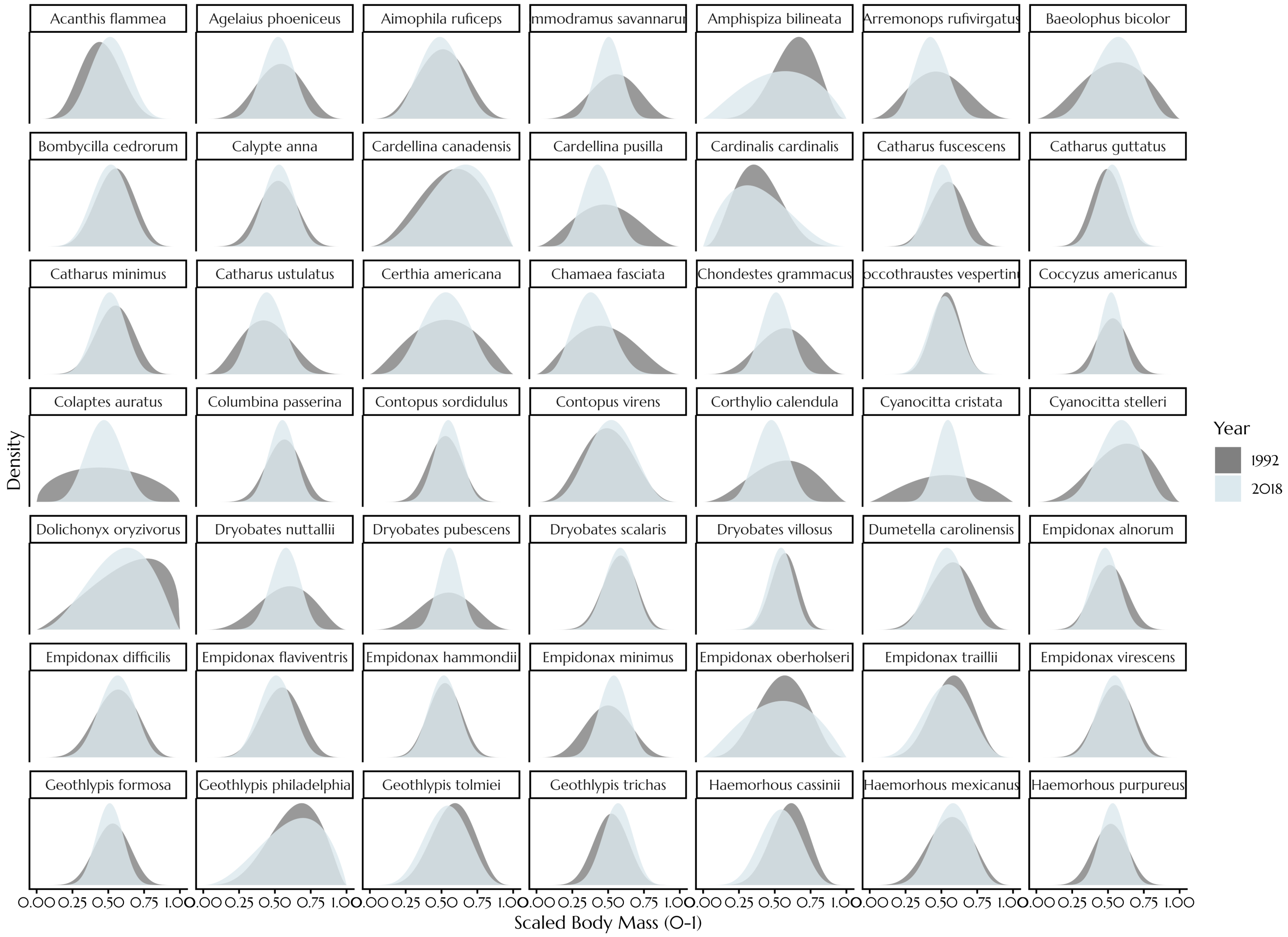

Acanthis flammea
Agelaius phoeniceus
Aimophila ruficeps
mmodramus savannaru
Amphispiza bilineata
Arremonops rufivirgatus
Baeolophus bicolor
Bombycilla cedrorum
Calypte anna
Cardellina canadensis
Cardellina pusilla
Cardinalis cardinalis
Catharus fuscescens
Catharus guttatus
Catharus minimus
Catharus ustulatus
Certhia americana
Chamaea fasciata
Chondestes grammacus
occothraustes vespertin
Coccyzus americanus
Colaptes auratus
Columbina passerina
Contopus sordidulus
Contopus virens
Corthylio calendula
Cyanocitta cristata
Cyanocitta stelleri
Dolichonyx oryzivorus
Dryobates nuttallii
Dryobates pubescens
Dryobates scalaris
Dryobates villosus
Dumetella carolinensis
Empidonax alnorum
Empidonax difficilis
Empidonax flaviventris
Empidonax hammondii
Empidonax minimus
Empidonax oberholseri
Empidonax traillii
Empidonax virescens
Geothlypis formosa
Geothlypis philadelphia
Geothlypis tolmiei
Geothlypis trichas
Haemorhous cassinii
Haemorhous mexicanus
Haemorhous purpureus
Density
0.00 0.25 0.50 0.75 1.00
Scaled Body Mass (0-1)
Year
1992
2018

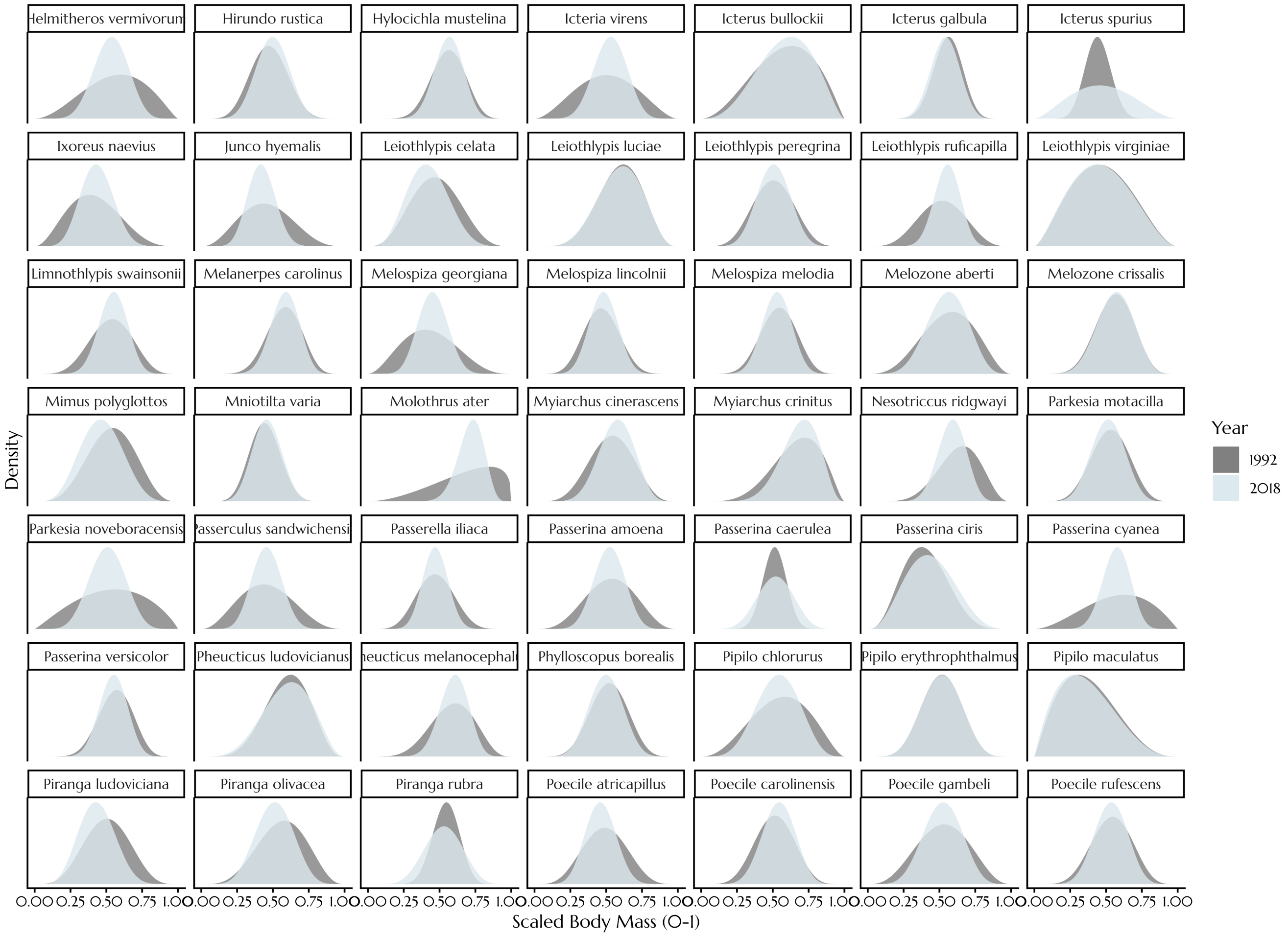

Helmitheros vermivorun
Hirundo rustica
Hylocichla mustelina
Icteria virens
Icterus bullockii
Icterus galbula
Icterus spurius
Ixoreus naevius
Junco hyemalis
Leiothlypis celata
Leiothlypis luciae
Leiothlypis peregrina
Leiothlypis ruficapilla
Leiothlypis virginiae
Limnothlypis swainsonii
Melanerpes carolinus
Melospiza georgiana
Melospiza lincolnii
Melospiza melodia
Melozone aberti
Melozone crissalis
Mimus polyglottos
Mniotilta varia
Molothrus ater
Myiarchus cinerascens
Myiarchus crinitus
Nesotriccus ridgwayi
Parkesia motacilla
Parkesia noveboracensis
asserculus sandwichensi
Passerella iliaca
Passerina amoena
Passerina caerulea
Passerina ciris
Passerina cyanea
Passerina versicolor
Pheucticus ludovicianus
heucticus melanocephal
Phylloscopus borealis
Pipilo chlorurus
Pipilo erythrophthalmus
Pipilo maculatus
Piranga ludoviciana
Piranga olivacea
Piranga rubra
Poecile atricapillus
Poecile carolinensis
Poecile gambeli
Poecile rufescens
Density
Year
1992
2018
0.00 0.25 0.50 0.75 1.00
Scaled Body Mass (0-1)

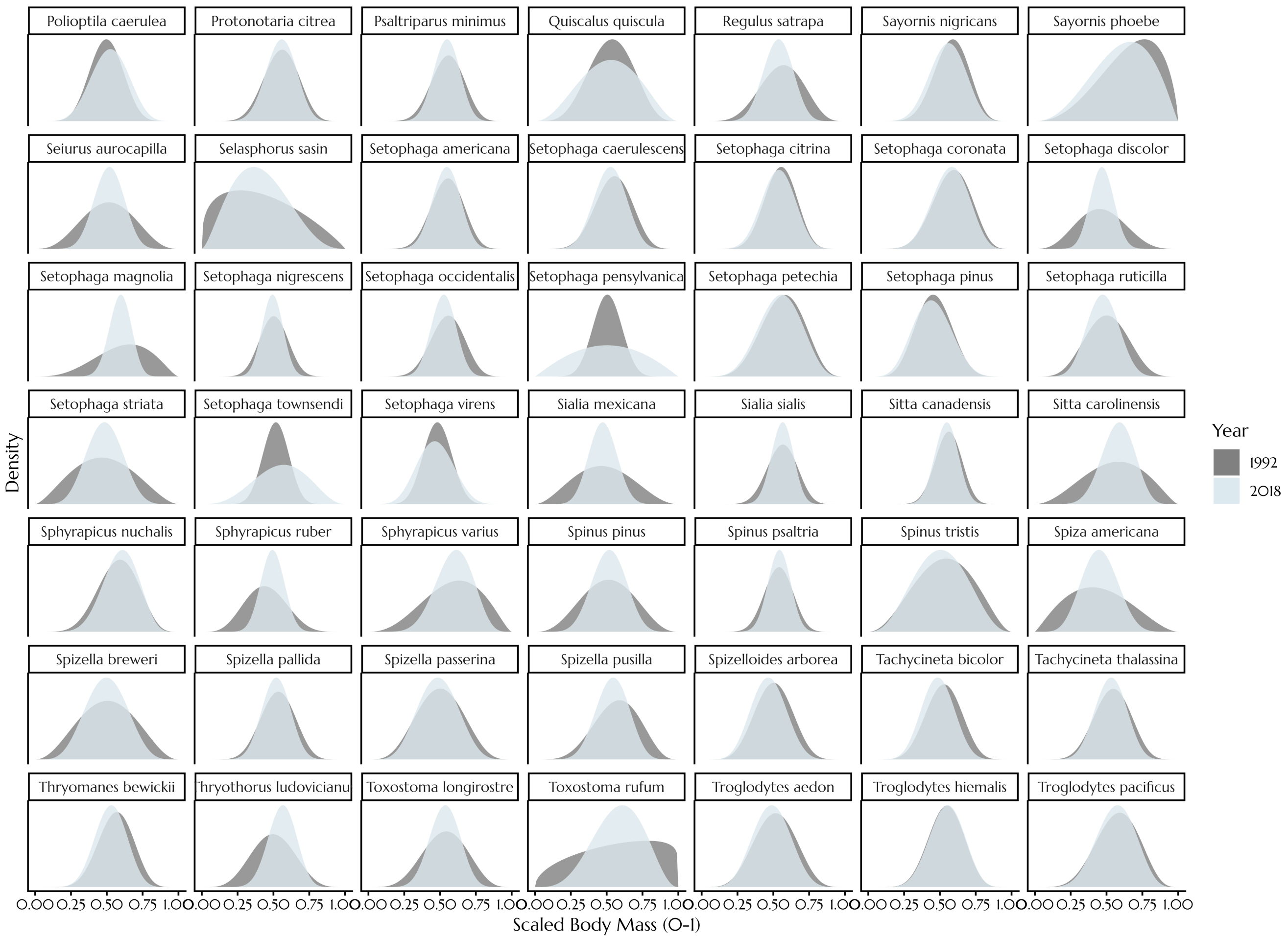
Polioptila caerulea
Protonotaria citrea
Psaltriparus minimus
Quiscalus quiscula
Regulus satrapa
Sayornis nigricans
Sayornis phoebe
Seiurus aurocapilla
Selasphorus sasin
Setophaga americana
Setophaga caerulescens
Setophaga citrina
Setophaga coronata
Setophaga discolor
Setophaga magnolia
Setophaga nigrescens
Setophaga occidentalis
Setophaga pensylvanica
Setophaga petechia
Setophaga pinus
Setophaga ruticilla
Setophaga striata
Setophaga townsendi
Setophaga virens
Sialia mexicana
Sialia sialis
Sitta canadensis
Sitta carolinensis
Sphyrapicus nuchalis
Sphyrapicus ruber
Sphyrapicus varius
Spinus pinus
Spinus psaltria
Spinus tristis
Spiza americana
Spizella breweri
Spizella pallida
Spizella passerina
Spizella pusilla
Spizelloides arborea
Tachycineta bicolor
Tachycineta thalassina
Thryomanes bewickii
Thryothorus ludovicianu
Toxostoma longirostre
Toxostoma rufum
Troglodytes aedon
Troglodytes hiemalis
Troglodytes pacificus
Density
Year
1992
2018
0.00 0.25 0.50 0.75 1.00
Scaled Body Mass (0-1)

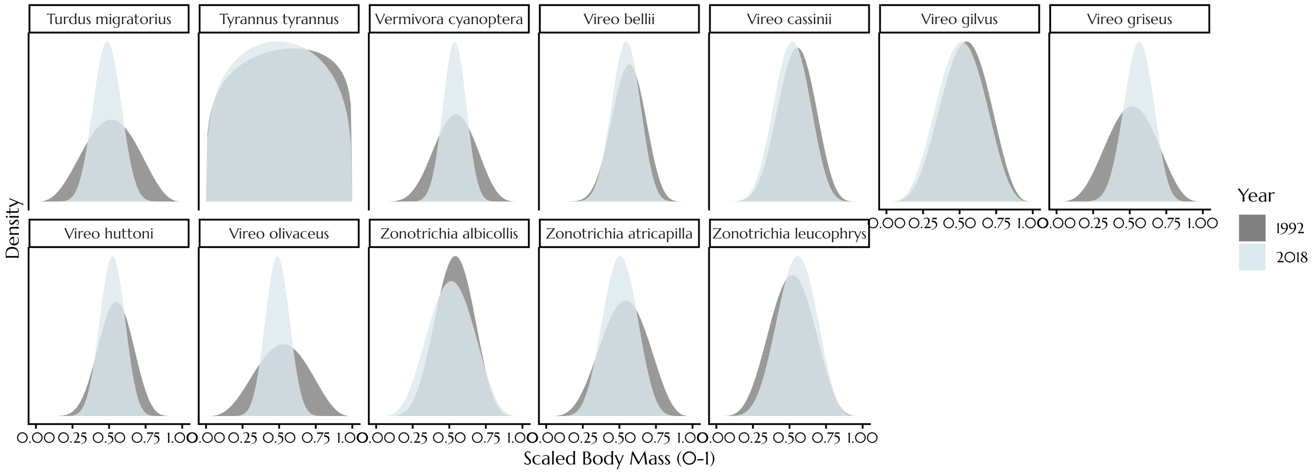


**Supplemental Figure 1 | Estimated body mass distributions of 159 North American bird species, between 1992 and 2018.** Distributions represent those at median observed range points per species, and were derived from a Bayesian beta-distributional model. Alpha and beta parameters used to draw each distribution represent posterior medians from our model.
